# Electrophysiological Activity of Primary Cortical Neuron-Glia Mixed Cultures

**DOI:** 10.1101/2022.12.29.522240

**Authors:** Noah Goshi, Hyehyun Kim, Gregory Girardi, Alexander Gardner, Erkin Seker

## Abstract

**Background:** We have previously described a tri-culture of neurons, astrocytes, and microglia that accurately mimics the *in vivo* neuroinflammatory response (both neurotoxic and neuroprotective) to a wide range of neuroinflammatory stimuli. Electrophysiological activity is an essential measure of neuronal function, which is influenced by neuroinflammation. Microelectrode array (MEA) technology is a versatile tool to non-invasively study *in vitro* neural networks by simultaneously recording electrophysiological activity from multiple sites. In this study, we used extracellular recordings to determine the effect of microglia on neural network formation and stability in primary cortical cultures and monitor the changes in neural activity in response to neuroinflammatory stimuli.

**Methods:** Primary neonatal rat cortical tri-cultures of neurons, astrocytes, and microglia or co-cultures of neurons and astrocytes were cultured on custom MEAs and the neural activity was monitored for 21 days *in vitro* to assess culture maturation and network formation. Quantification of synaptic puncta and averaged spike waveforms were used to determine the difference in excitatory to inhibitory neuron ratio (E/I ratio) of the neurons in tri- and co-cultures. The electrophysiological response to lipopolysaccharide (LPS) treatment of both culture types were compared.

**Results:** The tri- and co-culture showed minimal difference in electrophysiological markers of neural network formation and stability with the exception of a significant increase in spike frequency in the tri-culture at later timepoints (DIV 17 and 21). Additionally, there was no significant difference in the density of either post-synaptic or excitatory pre-synaptic puncta between the culture types. However, characterization of the average spike waveforms revealed that the tri-culture had an E/I ratio much closer to that found in the rat cortex. Finally, only the tri-culture displayed a significant decrease in both the number of active channels and spike frequency following LPS exposure.

**Conclusions:** This study demonstrates that the microglia in the tri-culture do not disrupt neural network formation and stability as quantified using extracellular recordings and may be a better representation of the *in vivo* cortex due to the closer E/I ratio than more traditional isolated neuron and neuron-astrocyte co-cultures. Additionally, the tri-culture is better able to mimic the neuroinflammatory response to LPS, which was quantified via changes in neural electrophysiological activity.

## Introduction

*In vitro* models of the central nervous system (CNS) are powerful tools that allow researchers to conduct highly directed cellular level experiments to study the cellular and molecular pathways underlying many neurological disorders. However, there are inherent limitations within these models that limit their physiological relevance, most notably researchers much ensure that the appropriate cell types are included within their culture model to effectively recapitulate the *in vivo* disease state. This is especially true for CNS models as crosstalk between neurons and glia play significant roles in both pathological conditions [1–5] and maintaining homeostasis [6,7]. Microelectrode array (MEA) technology is a popular method to study *in vitro* neural networks, as it provides a non-invasive method to simultaneously record electrophysiological activity from multiple sites [8,9]. This allows researchers to conduct longitudinal studies to study neural network formation [10,11] and disruption caused by neurotoxic compounds [12–14] and proteins associated with neurodegeneration [15–17]. Nevertheless, the physiological relevance of these studies is still largely dependent on the cell types present within the culture and may not fully recapitulate the *in vivo* response [18]. Therefore, there has been significant effort in developing new CNS culture models and characterizing the spatial and temporal aspects of the electrophysiological recordings both during spontaneous activity and in response to stimuli. This includes studying the neural activity from cultures of neurons from different regions of the brain (cortex [8,19,20] vs hippocampus [21,22]), neurons cultured with supporting glial cells such as astrocytes [23] and oligodendrocytes [24], and human primary [25] or stem cell derived [26,27] neurons. Additionally, electrophysiological recording from complex *in vitro* models such as 3D [28,29] and organ-on-a-chip [30,31] models have been studied to further improve the physiological relevance of *in vitro* neural models.

We have previously developed a neuron, astrocyte, and microglia tri-culture to study neuroinflammation *in vitro* [32]. This tri-culture is maintained by culturing primary cortical neurons from neonatal rats in a serum-free culture medium specifically designed to support all three cell types. We have demonstrated that the tri-culture model better mimics the neuroinflammatory response to a number of stimuli including lipopolysaccharide (LPS) exposure, mechanical trauma, glutamate induced excitotoxicity, and toll-like receptor (TLR) activation [32,33] as compared to neuron-astrocyte co-cultures. In this paper we further explore the capabilities of the tri-culture by characterizing the neural functionality of the system. We compared the extracellular recordings taken over 21 days *in vitro* (DIV) between the tri- and co-cultures to determine the effect of microglia on network formation and neuron function. We demonstrate that many features of neural activity remain similar between the tri- and co-cultures (% active channels, spike frequency, and burst characteristics), suggesting that the presence of microglia do not adversely affect the neuronal function. The tri-culture shows increased synchrony at early timepoints compared to the co-culture, which may indicate faster maturation of network activity in the tri-culture configuration. Additionally, analysis of the action potential waveform characteristics reveals that the tri-cultures contain an increased excitatory/inhibitory (E/I) neuron ratio that more closely resembles the ratio found *in vivo*, and this observation is supported by synaptic staining. We also incorporated the tri-culture into compartmentalized organ-on-a-chip devices that separate the neural axons and somas using microchannels [31,34], and recorded robust neural activity from these platforms. Finally, we demonstrate the ability to observe changes in neural activity in response to known inflammatory stimuli (LPS exposure).

## Methods

### MEA design and fabrication

Custom MEAs and microfluidic platforms were fabricated using previously described methods [31]. Briefly, standard (well) MEAs were designed with a 4 x 8 array of electrodes (32 total) each with a diameter (Ø) of 20 μm and an interelectrode pitch of 130 μm. The electrodes and traces (250 nm-thick Au over a 160 nm-thick Cr adhesion layer) were sputter-deposited on borosilicate glass wafers (500 μm thick, University Wafers) and patterned using standard lift-off techniques). SiO_2_ was deposited via PECVD to serve as the insulation layer. Finally, the electrode sites were lithographically patterned and opened via a brief immersion in buffered oxide etch. Glass cloning cylinders (8 mm x 6 mm inner Ø, Sigma) were then attached over the MEA using sterile vacuum grease (Dow Corning).

Microfluidic platforms were designed following recommendations from our previous study [31]. Polydimethylsiloxane (PDMS; Slygard 184, Dow Corning) microfluidic devices were fabricated using standard soft-lithography techniques. These devices contained two large cell culture chambers connected by 101 microchannels with dimensions of 1000 μm x 10 μm x 1.5 μm (length x width x height). These microchannels allowed axons, but not somas of neurons to enter the microchannels and thereby synaptically connect the two separate neural populations. Corresponding MEAs with an array of 8 microchannels in each cell culture chamber and an array of 16 electrodes placed under the microchannels were fabricated using the same methods as above. To form the final device, both the MEA and PDMS platforms were sterilized with 70% EtOH and the surfaces were activated with air plasma at 10 W for 2 min. The MEA was then covered with 70% EtOH and the PDMS platform was placed over the MEA and aligned under a microscope. The aligned device was placed in a vacuum chamber for 1 h to remove the EtOH solution and permanently bond the MEA and PDMS platform. The bonded devices were then treated with air plasma at 30 W for 10 min to make the surfaces hydrophilic and glass cloning cylinders were mounted over the fluidic ports.

### Primary cortical culture

All media were prepared as previously described [19,32]. Briefly, plating medium consisted of Neurobasal A culture medium supplemented with 2% B27 supplement, 1x GlutaMAX, 10% heat-inactivated horse serum, and 20 mM HEPES at pH 7.5, while the co-culture medium consisted of Neurobasal A culture medium supplemented with 2% B27 supplement and 1x GlutaMAX (all from ThermoFisher). The tri-culture medium consisted of supplementing the co-culture medium with 100 ng/mL mouse IL-34 (R&D Systems), 2 ng/mL TGF-β (Peprotech), and 1.5 μg/mL ovine wool cholesterol (Avanti Polar Lipids), which were identified as factors that support isolated microglia survival in culture [35]. Due to the limited shelf life of IL-34 and TGF-β, the tri-culture medium was made fresh each week.

All procedures involving animals were conducted in accordance with the National Institutes of Health Guide for the Care and Use of Laboratory Animals following protocols approved by the University of California, Davis Institutional Animal Care and Use Committee. Timed-pregnant Sprague-Dawley rats were purchased from Charles River Laboratory (Hollister, CA). All animals were housed in clear plastic shoebox cages containing corn cob bedding under constant temperature (22 ± 2 °C) and 12 h light-dark cycle. Food and water were provided *ad libitum*. Primary cortical cultures were prepared from postnatal day 0 rat pups as previously described [36]. Neocortices from all pups in the litter were pooled, dissociated, and resuspended in plating medium. Prior to plating all substrates were coated with 0.5 mg/mL poly-L-lysine (Sigma) in B-buffer (3.1 mg/mL boric acid and 4.75 mg/mL borax, Sigma) for 4 h at 37 °C and 5% CO_2_, and then washed with sterile deionized water and covered with plating medium. Cultures were plated at a density of 550 cells/mm^2^ on well MEAs (**Figure S1**), while for the microfluidic devices 20 μL of cell suspension at a concentration of 3×10^6^ cells/mL were seeded in each chamber (**Figure S2**). The cells were allowed to adhere for 4 h, after which the plating medium was changed to tri- or co-culture medium. Half-media changes were performed every 3-4 days with the respective media types.

### Immunocytochemistry

At the conclusion of the experiment, the cell cultures were washed 3 times with 37 °C DPBS+ and fixed using 4% w/v paraformaldehyde (PFA; Affymetrix) in PBS for 2.5 h. Fixed cells were washed twice with 0.05% v/v Tween20 (Sigma) solution in DPBS+, followed by a 3 min permeabilization with 0.1% v/v Triton X-100 (ThermoFisher) solution in DPBS+ and two additional washes with Tween20 solution. Samples were blocked with a solution of 0.5% v/v heat-inactivated goat serum (ThermoFisher) and 0.3 M glycine (Sigma) in DPBS+ (blocking buffer) for 1 h. Following the blocking step, samples were incubated for 1 h in primary antibody solution containing mouse anti-PSD-95 (ThermoFisher), rabbit anti-VGlut1 (ThermoFisher), and chicken anti-MAP2 (Abcam) in blocking buffer. Alternatively, some samples were incubated with mouse anti-βIII tubulin (ThermoFisher), rabbit anti-GFAP (ThermoFisher), and chicken anti-Iba1 (Abcam) to ensure the cellular compositions of the culture. Samples were then washed 3 times with Tween20 solution before a 1 h incubation with secondary antibody solution containing goat anti-mouse antibodies conjugated to AlexaFluor 647 (ThermoFisher), goat anti-rabbit antibodies conjugated to AlexaFluor 488 (ThermoFisher), and goat anti-chicken antibodies conjugated to AlexaFluor 555 (ThermoFisher). Following incubation with secondary antibody solution, the samples were washed 3 times with DPBS+. Lastly, samples were incubated for 5 min with a 4’,6-diamidino-2-phenylindole (DAPI) solution (Sigma) to stain cell nuclei, followed by an additional Tween20 solution wash before mounting them onto glass slides using ProLong Gold Antifade Mountant (ThermoFisher).

### Image analysis

All sample images were acquired using a Leica TCS SP8 STED 3X microscope with a 63x/1.4 oil immersion objective and analyzed using a custom ImageJ macro [32]. Briefly, the images were thresholded to determine the number of pre- and post-synaptic puncta, and the number of mature synapses were quantified by determining the number of co-localized puncta. These values were then correlated with dendrite length to determine the number of puncta or mature synapses per 100 μm of dendrite length. For all analysis five predetermined fields were analyzed per well to account for variability within the individual cultures.

### Electrophysiological recording and analysis

For extracellular electrophysiological recordings, the devices were placed on a custom-built rig and maintained at 37 °C and 5% CO_2_ during the recordings. Recordings were performed at a sampling frequency of 30 kHz using an RHD2132 Intan amplifier (Intan Technologies). Half-media changes were performed 24 h prior to each recording except for LPS treated conditions, in which the half-media change was performed 24 h prior to the baseline measurement.

Feature extraction from the recordings was done using Offline Sorter and NeuroExplorer (Plexon). Spikes were detected following high-pass filtering (300 Hz cut-off) using a threshold of ±8x standard deviation of the noise. Channels that showed less than 10 spikes per 10 min recording were discarded from the analysis. The number of active electrodes for each array was determined as the number of electrodes that registered at least 10 spikes during the 10 min recording. Overall synchrony of the active electrodes within each device was determined using the SPIKE-distance methodology [37] in the PySpike Python package [38]. Bursts were classified using the *max interval* method in NeuroExplorer using previously defined parameters [24,39]. Briefly these parameters were; (i) maximum initial inter-spike interval (ISI) of 0.1 s, (ii) maximum end ISI of 0.25 s, (iii) minimum inter-burst interval of 0.5 s, (iv) minimum burst duration of 0.05 s, and (v) minimum number of spikes in bursts of 6 spikes, and were used to determine the percentage of spikes in bursts, average burst duration, and average interspike interval (ISI) within bursts at a per electrode basis. Spike frequency and burst features were determined on a per electrode basis, but statistical analysis was performed on a per device basis by taking the overall mean from the active electrodes.

To determine the average peak-trough duration, we first sorted the extracted spikes using the valley-seeking algorithm in OfflineSorter (Plexon) with a Parzen multiplier of 2.0. Due to the relatively large distance between the electrodes and low seeding density, nearly every electrode only displayed one cluster; however, on occasion two duplicate clusters were found due to the use of dual thresholds during spike detection (±8σ). As these two clusters clearly belonged to the same group, one cluster was disregarded from the analysis. The average peak-trough duration of each cluster was then calculated via OfflineSorter.

### Statistical analysis

To compare change in the different spike features based on culture type or condition, each data set was fitted using a linear mixed effects model (treating the individual devices as a random effect) with a b-spline basis to account for the overall shape of the plots [31]. We then compared the estimated marginal means from the fitted curves at each timepoint and adjusted for multiple comparisons using the Holm-Bonferroni method. A 2-way ANOVA was used to compare the differences in synaptic staining overtime, while a Student’s t-test was used to compare the differences in peak-trough duration between the tri- and co-culture at DIV 21. For all experiments, statistical significance was determined by *p*-values < 0.05, and results are given as mean ± standard error of the mean (SEM).

## Results

### Electrophysiology

We recorded spontaneous neural activity from DIV 7 to DIV 21 from both tri- and co-cultures to determine if there was any discernible difference in culture maturation or network formation (**Figure 1a,b**). As expected, we observed a significant increase in the percentage of active channels in both the tri- and co-culture from DIV 7 to DIV 21 (**Figure 1c**). However, we did not find a significant difference in the percentage of active channels between the tri- and co-cultures at any timepoint. We saw a similar trend in spike frequency for both the tri- and co-cultures (**Figure 1d**), with no differences between the tri- and co-cultures. However, in this case the increase in spike frequency was more subtle, and therefore we did not observe a significant increase between DIV 7 and later timepoints with the exception of the tri-culture at DIV 21 (*p* = 0.0076). We also compared other electrophysiological features used to determine culture maturation and stability over time (**Figure S3**), such as the percent of spikes in bursts, burst duration, and ISI within bursts. Both the tri- and co-cultures showed a significant increase in the percentage of spikes in bursts indicating that the cultures were maturing as expected; however, there was no difference between the two cultures (**Figure S3a**). Similarly, we saw no difference between the average burst duration and within burst ISI between the two culture types (**Figure S3b,c**), and for both the co- and tri-culture these values remained stable over time, suggesting no degradation in neural health over the 21 DIV window. Finally, we compared network formation in the tri- and co-cultures by assessing the synchrony of the spike trains from the active electrodes in each culture using the SPIKE-distance method [37,38]. It is important to note that the SPIKE distance method is a measurement of spike-train dissimilarity given on a scale of 0-1. Therefore, in order to measure the synchrony of the culture, we subtracted the SPIKE distance value from 1, with values closer to 1 indicating increased synchrony in the culture. We observed a significant increase in synchrony over time in both the co- and tri-cultures indicating robust network formation (**Figure 1e**). While there was no significant difference in the synchrony between the tri- and co-culture, there was a trend towards increased synchrony in the tri-culture at DIV 10 (*p* = 0.11).

**Figure 1:**
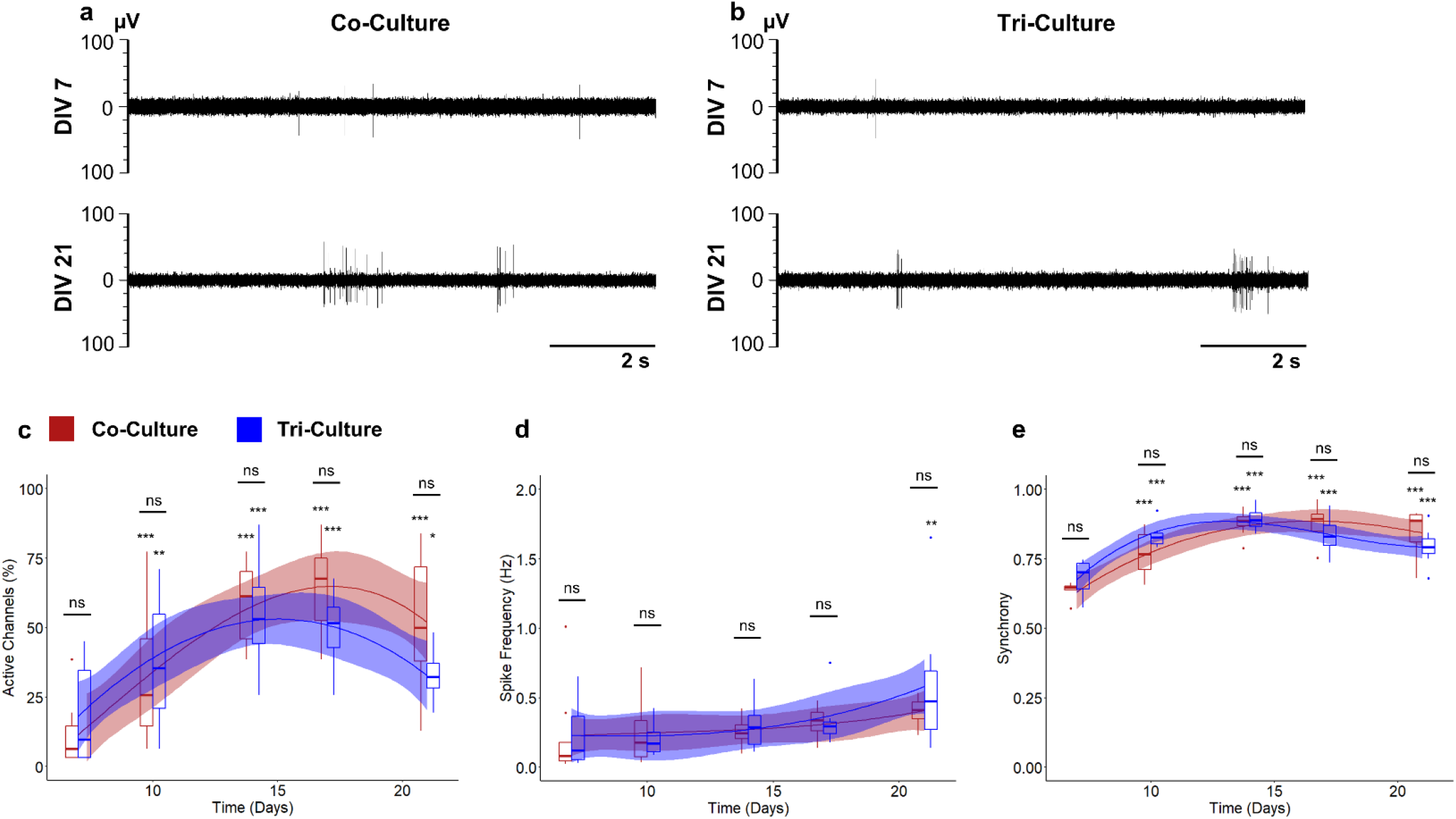
Representative extracellular recordings taken at DIV 7 and DIV 21 from (a) co-cultures and (b) tri-cultures. Comparisons of the (c) percent active channels, (d) spike frequency, and (e) synchrony between co-cultures (red) and tri-cultures (blue). The solid lines show the fitted linear mixed effects model (treating individual cultures as a random effect) with a b-spline basis. The shaded regions are the 95% confidence interval. An asterisk above an individual box indicates a significant difference of the estimated marginal means of the fitted curves between that timepoint and DIV 7 of the same culture type, while the bars indicate the significance between the co- and tri-culture at that timepoint (*n* = 8, from three independent dissections). \**p* < 0.05, \*\**p* < 0.01, \*\*\**p* < 0.001, ns indicates no significant difference.

### Incorporation into microfluidic platforms

In order to further investigate some of the non-statistically significant, but important trends we previously observed, we compared the spontaneous neural activity from tri- and co-cultures maintained in microfluidic devices, which have been previously shown to improve electrophysiological recording fidelity [31,40,41]. We once again demonstrate that both the tri- and co-cultures show significant increase in the percentage of active channels over time with no significant difference between the cultures (**Figure 2a**). However, using the microfluidic device, we found that both the tri- and co-cultures show significant increases in spike frequency from axons within the microchannels over time, and we observed significantly higher spike frequencies in the tri-culture at DIV 17 and 21 (*p* = 0.016 and *p* = 0.0017 respectively; **Figure 2b**). Additionally, we observed no differences among other electrophysiological features (percent spikes in bursts, burst duration, and within burst ISI; **Figure S4**) and spike train synchrony (**Figure 2c**) between the tri- and co-cultures, once again suggesting minimal differences in culture maturation and stability between the tri- and co-cultures, with the exception of increased spike frequency in the tri-culture.

**Figure 2:**
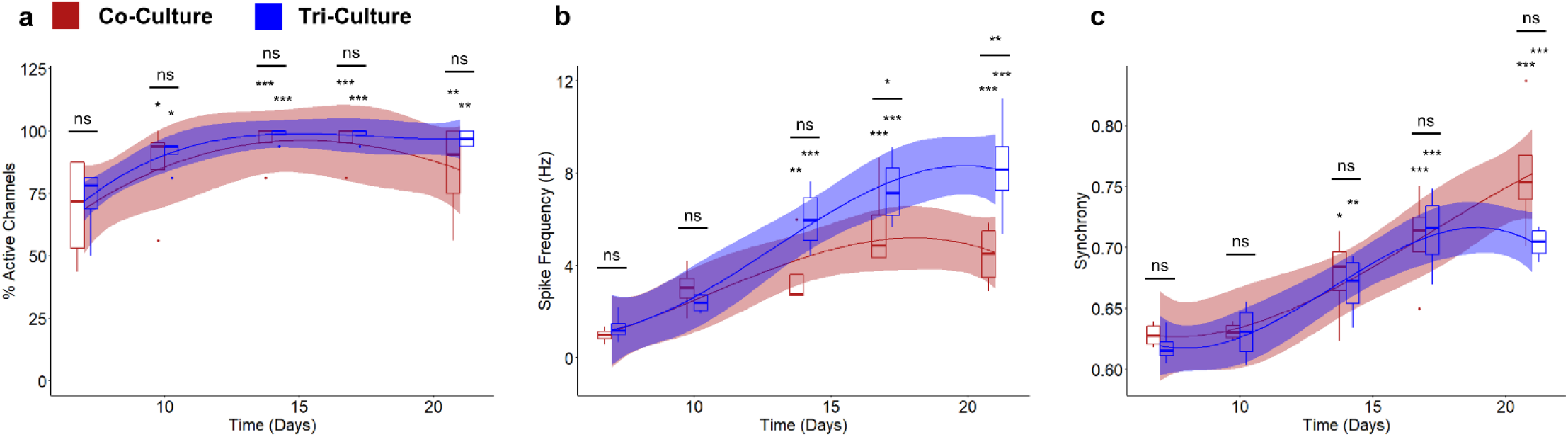
Comparisons of the (a) percent active channels, (b) spike frequency, and (c) synchrony between co-cultures (red) and tri-cultures (blue) cultured in a two-chambered microfluidic device. The solid lines show the fitted linear mixed effects model (treating individual cultures as a random effect) with a b-spline basis. The shaded regions are the 95% confidence interval. An asterisk above an individual box indicates a significant difference of the estimated marginal means of the fitted curves between that timepoint and DIV 7 of the same culture type, while the bars indicate the significance between the co- and tri-culture at that timepoint (*n* = 5, from two independent dissections). \**p* < 0.05, \*\**p* < 0.01, \*\*\**p* < 0.001, ns indicates no significant difference.

### Synaptic density

As microglia are known to play a significant role in synapse formation and elimination during postnatal development [42–44], we compared synapse formation between the tri- and co-cultures over 21 DIV (**Figure 3a**). A two-way ANOVA revealed that there was no significant interaction between culture type and time in culture on the density of post-synaptic marker PSD-95 (**Figure 3b**) and excitatory pre-synaptic marker VGlut1 (**Figure 3c**) puncta (p = 0.46 and p = 0.36 respectively). Simple main effects analysis revealed no significant difference between the density of PSD-95 and Vglut1 puncta based on culture (*p* = 0.26 and *p* = 0.15 respectively) but did reveal a significant difference based on time in culture (*p* = 0.0032 and *p* = 00.15 respectively). We also compared the number of co-localized puncta as a measurement of mature excitatory synapses [45,46] (**Figure 3d**). Once again, a two-way ANOVA revealed no-significant interaction between culture type and time in culture on the number of mature excitatory synapses (*p* = 0.22), and simple main effects analysis revealed a significant difference increase in mature excitatory synapses based on time in culture (*p* = 0.0066) but not based on culture type (*p* = 0.21).

**Figure 3:**
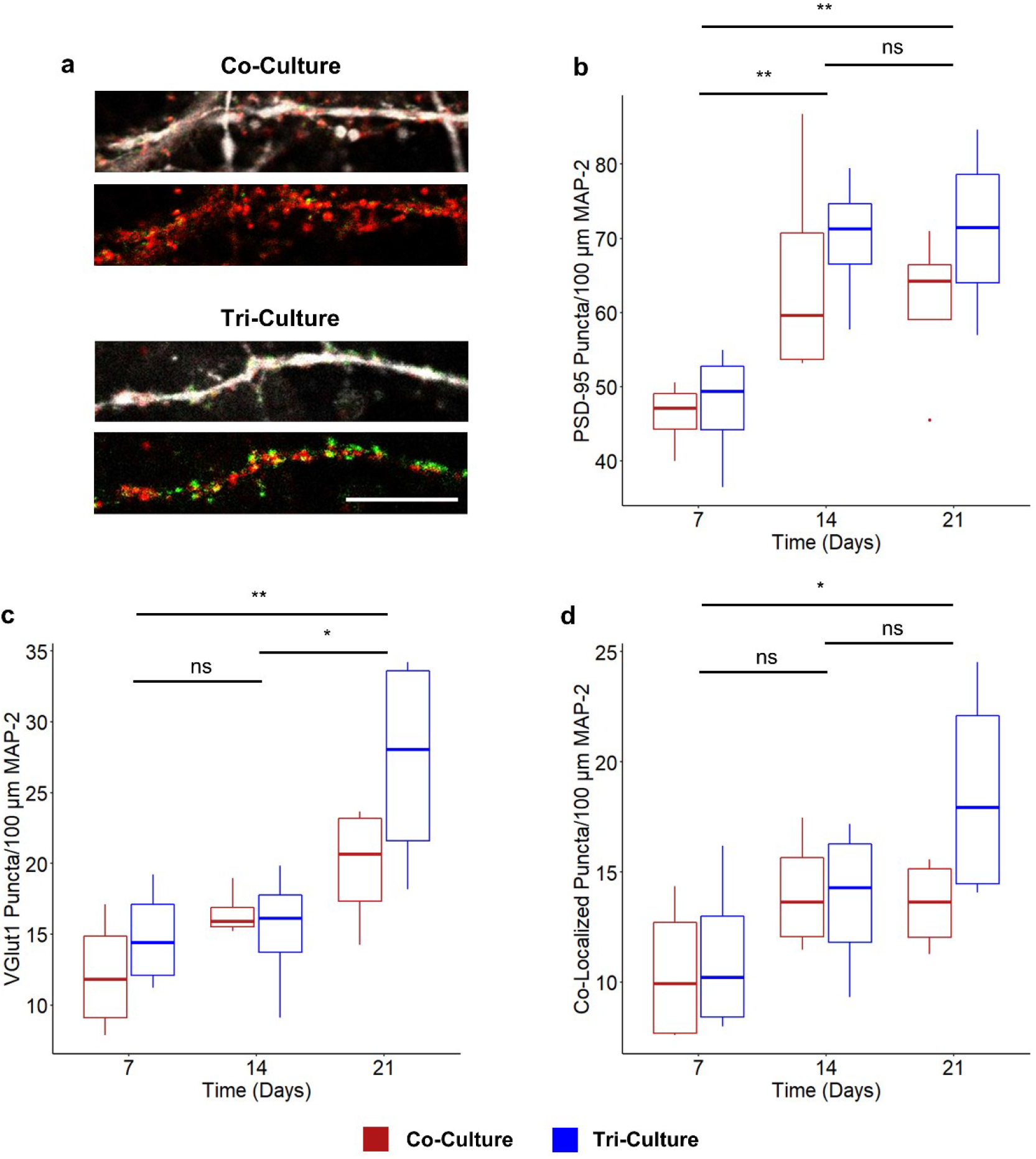
(a) Representative fluorescence images of co- and tri-cultures at DIV 21. The cultures are stained for the excitatory pre-synaptic marker VGlut1 (Green), post-synaptic marker PSD-95 (red). The above image also shows the co-localization with MAP-2 (white). (scale bar = 10 μm). Comparison of the density of (b) PSD-95 puncta, (c) VGlut1 puncta, and (d) co-localized puncta. In all three cases a two-way ANOVA found no interaction between culture type and time in culture, and therefore the asterisk indicates the significance of the main effect between timepoints (*n* = 4, from two independent dissections). \**p* < 0.05, \*\**p* < 0.01, ns indicates no significant difference.

### Excitatory vs. Inhibitory Neurons

While we did not observe any statistically-significant differences in synapse formation between the tri- and co-culture (**Figure 3d**), we saw evidence of increased excitatory synapse formation at DIV 21 compared to DIV 7 in the tri-culture (**Figure 3d**) and a significant increase in spike frequency in the tri-culture (**Figure 2b**). As the cortex contains both excitatory and inhibitory neurons, we asked if the aforementioned differences could be attributed to changes in the ratio of excitatory to inhibitory neurons (E/I ratio). We found a bimodal distribution of the peak-trough duration of the averaged spike waveforms from 249 units recorded from both tri- and co-cultures (**Figure S5**). The first peak is centered at a peak-trough duration of ~220 μs, which correlates well with narrow-spiking inhibitory interneurons, while the second peak is centered at a peak-trough duration of ~380 μs, which correlated with broad-spiking excitatory neurons [47–49]. Additionally, the average spike waveforms from these two groups correlated well with previously described narrow-spiking and broad-spiking waveforms (**Figure 4a**) [48,49]. Using the density histogram as a guide, we classified units as either inhibitory (peak-trough duration < 280 μs) or excitatory (peak-trough duration > 280 μs). We observed a significant increase in E/I ratio of recorded units in the tri-culture (86.91 ± 3.27% excitatory neurons) as compared to the co-culture (68.83 ± 5.55% excitatory neurons; *p* = 0.016), as shown in **Figure 4b**. Interestingly, E/I ratio of the tri-culture more closely resembles the E/I ratio of the cortex *in vivo* (~80% excitatory) [50].

**Figure 4:**
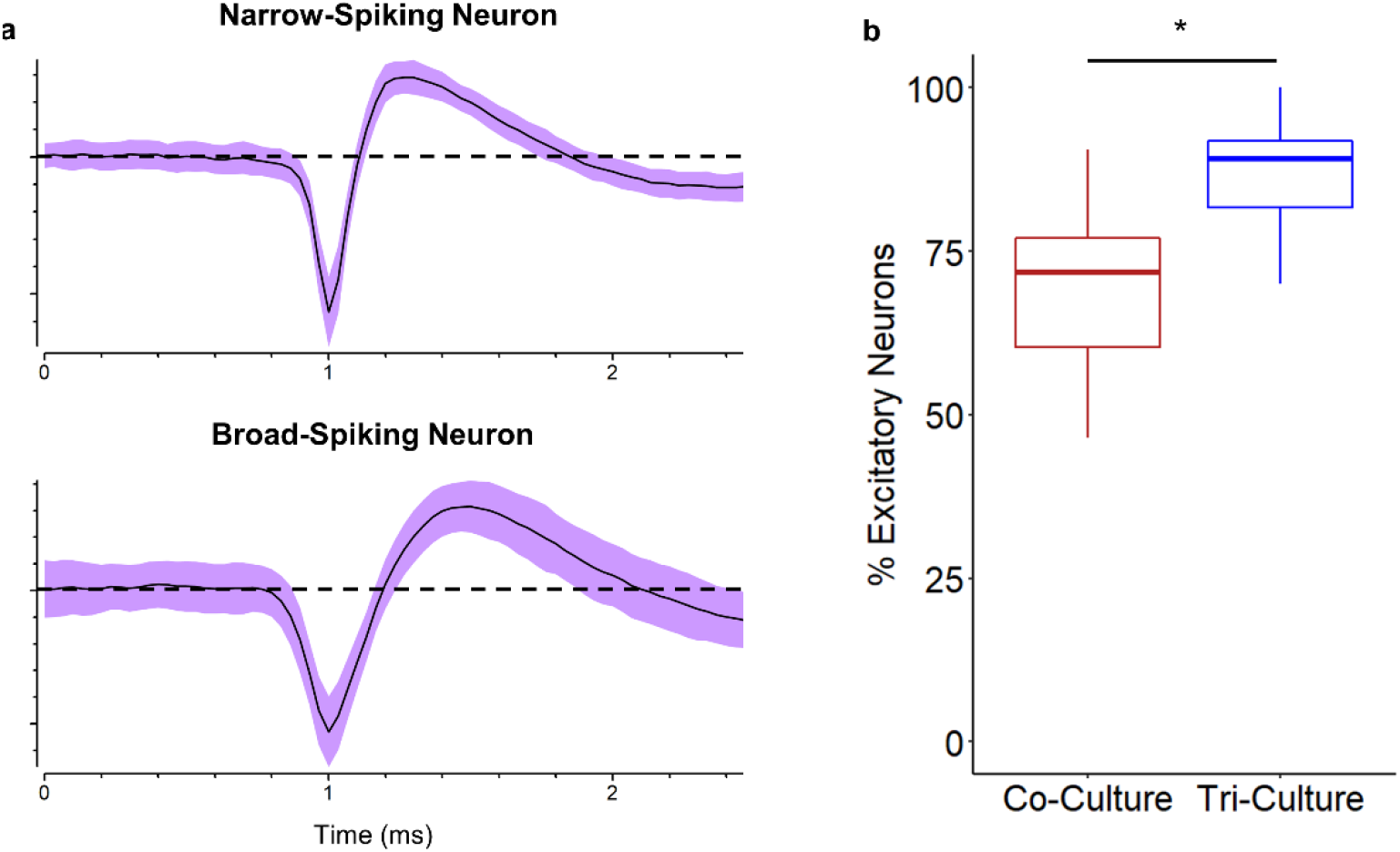
Comparison of the E/I ratio in tri- and co-cultures at DIV 21. (a) representative waveforms of narrow-spiking and broad-spiking neurons recorded at DIV 21. The line represents the average spike waveform, while the shaded region is one standard deviation. (b) Percent excitatory neurons in the tri- and co-cultures at DIV 21 (n = 8 MEAs (249 total units), from three independent dissections). \**p* < 0.05.

### Influence of LPS Exposure

In order to demonstrate the ability to detect changes in neural activity in response to neuroinflammatory stimuli, we challenged both co- and tri-cultures with 5 μg/mL LPS at DIV 21. As expected, there was no change in the neural activity in the co-culture in response to LPS, as LPS acts through the toll-like receptor 4 (TLR4), which is primarily expressed by microglia. Conversely, we observe a significant decrease in both active channels (**Figure 5a**) and spike frequency (**Figure 5b**) in the tri-culture following exposure to LPS. Additionally, we begin to see a trend towards a reduction in spike frequency beginning at 3 h (*p* = 0.17) before there is any change in the number of active channels (*p* = 0.71). We were unable to compare other electrophysiological features (percent spikes in bursts, burst duration, and within burst ISI) and spike train synchrony, due to the dramatic reduction in active channels in the tri-culture following exposure to LPS limiting our ability to effectively characterize these features.

**Figure 5:**
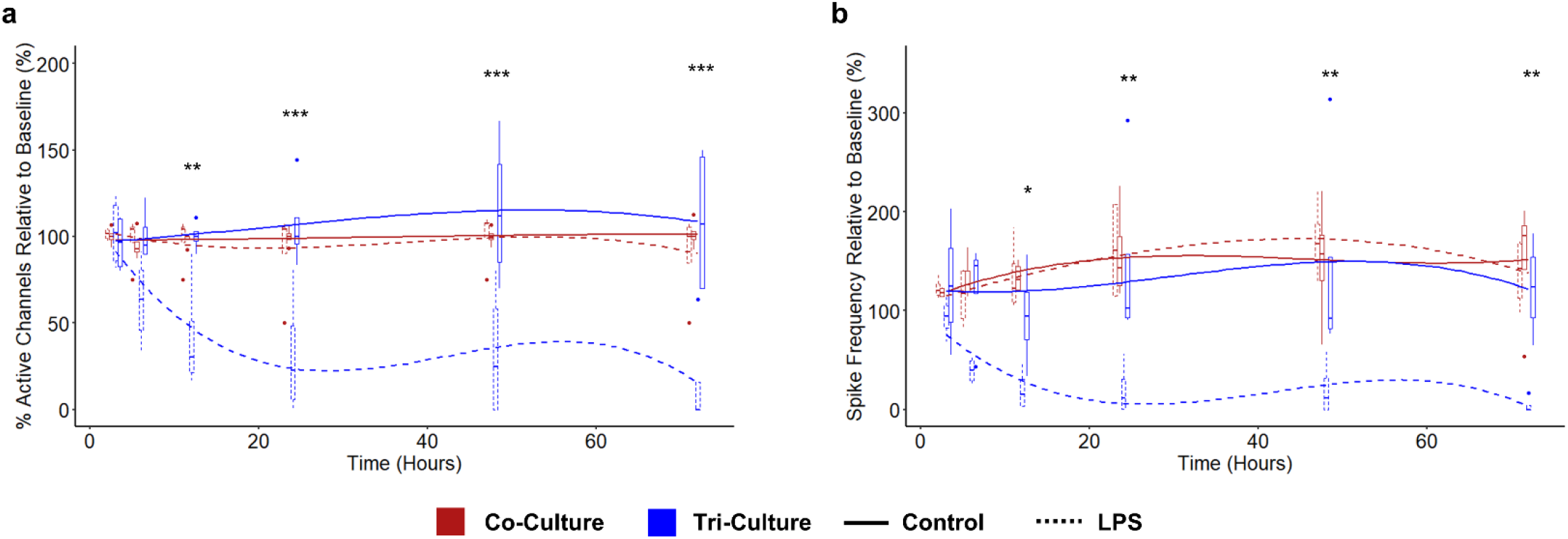
Comparing the change in (a) percent active channels and (b) spike frequency following exposure to 5 μg/mL LPS between co-cultures (red) and tri-cultures (blue). The lines show the fitted liner mixed effects model (treating individual cultures as a random effect) with a b-spline basis. Asterisks indicate a significant difference in the estimated marginal means of the fitted curves between control and LPS treated tri-cultures at that timepoint (*n* = 4, from two independent dissections). \**p* < 0.05, \*\**p* < 0.01, \*\*\**p* < 0.001.

## Discussion

Extracellular recording techniques using MEAs are a powerful tool to study *in vitro* neural cultures as it provides a non-invasive method to simultaneously record neural activity from multiple neurons within a culture. This makes it a popular method to study the changes in neural activity over time, as multiple recordings can be taken from the same culture to monitor changes during neural network formation [10,11] or in response to neurotoxic compounds [12–14]. In this study, we have characterized the neural network maturation and electrophysiological response to neuroinflammatory stimuli of a recently described tri-culture of neurons, astrocytes, and microglia that has been shown to better mimic the *in vivo* neuroinflammatory response [32,33,51]. Additionally, we compared the neural activity of the tri-culture to a well characterized neuron-astrocyte co-culture [23] to not only ensure that the presence of microglia in the tri-culture do not negatively impact the overall health of the neurons, but also to determine if they may influence neural activity or network formation as microglia are known to play a significant role in synapse formation and plasticity [44,52,53]. As expected, we observed many of the common indicators of neural network formation and maturation including an increase in the number of active channels, increased spike frequency, and increased synchrony in both the tri- and co-culture over time (**Figure 1**), indicating that the microglia within the tri-culture did not disrupt culture maturation or health. Similarly, we observed no significant differences between the tri- and co-cultures when comparing any of the aforementioned indicators of culture maturation (**Figure 1**) or other electrophysiological features (**Figure S1**) at any timepoint, which was somewhat surprising considering the significant role microglia are known to play in synaptic plasticity. However, we did see a trend towards increased neuronal network maturation in the tri-culture, as the synchrony of the tri-culture was increased at earlier timepoints (DIV 10, *p* = 0.11), but this increase was not statistically significant (**Figure 1e**). Additionally, we saw some evidence that the tri-culture had an increase in spike frequency at DIV 21 as only the tri-culture showed a significant increase in spike frequency between DIV 7 and DIV 21 (*p* = 0.0076 vs *p* = 0.51 for the co-culture), but once again there was no significant difference in spike frequency between the tri- and co-cultures at DIV 21 (**Figure 1d**). A recent study using a similar tri-culture showed a reversed trend, with microglia appearing to reduce spike frequency and other electrophysiological features [54]. However, in that paper, the amount of microglia was increased to 15-25% of the total cell population, which is more than double the number of microglia found in our tri-culture model [32] and the proportions found *in vivo* [55]. Furthermore, the authors found that increased microglia reactivity with increasing microglia density, suggesting that the decrease in spike frequency may be a function of reactive microglia as opposed to more homeostatic microglia and would be in line with the results we obtained from our LPS treated cultures.

In order to further investigate the potential difference in spike frequency, we cultured both the tri- and co-cultures in microfluidic devices, which have previously been shown to improve electrophysiological recording fidelity [31]. These devices consist of two cell culture chambers connected by a series of small microchannels with an integrated MEA. The small cross-sectional area of the microchannels prevent the cell bodies of neurons and glia from entering the microchannels, but permit axons to pass through and synaptically connect the two neural cultures. Additionally, the spatial confinement forces axons in close approximation to electrodes placed underneath the microchannels and the reduced volume within the microchannels reduces ionic dispersion leading to an increase in spike amplitude [56], which leads to an overall improvement in recording fidelity. This enhancement in recording fidelity allowed us to not only observe the expected increase in spike frequency in both the tri- and co-cultures indicative of culture maturation [57,58], but also revealed a significant increase in the spike frequency of the tri-culture at later timepoints (**Figure 2b**), which is in line with the observations from the recordings taken from standard MEAs (**Figure 1d**). Furthermore, incorporation of the tri-culture into the microfluidic device was straightforward as it only required the use of a specialized culture medium, highlighting the tri-culture’s amenability to complex culture setups such as microfluidic devices and organ-on-a-chip platforms.

In order to determine the cause of increased spike frequency in the tri-culture at later timepoints, we compared the number of pre-synaptic and excitatory post-synaptic puncta in the tri- and co-cultures, as microglia are known to play a significant role in synaptic formation and elimination during postnatal development [43,44]. Interestingly, a 2-way ANOVA did not establish an interaction between time in culture and culture type (tri-vs co-culture) when comparing the density of pre-synaptic, excitatory post-synaptic, or co-localized puncta (**Figure 3b-d**). Additionally, main effects analysis did not reveal a significant difference between the tri- and co-cultures for any of the conditions. This was an unexpected finding as previous *in vitro* studies have shown that IL-10 secreted from microglia increase spine density and excitatory synapses [59]. However, in that study the microglia and neurons were physically separated, with the microglia grown on a porous insert that was added to the isolated neuron culture at a later timepoint, preventing their physical contact, and it has been shown that direct microglia contact with synapses can lead to synaptic elimination [60,61]. While we did not observe a significant difference between the tri- and co-culture, we noted a trend towards increased excitatory post-synaptic puncta at DIV 21 in the tri-culture (**Figure 3c**). Additionally, we observe that the tri-culture continues to show increased co-localized puncta over the 21 DIV, while the number of co-localized puncta appears to plateau at DIV 14 (**Figure 3d**). The fact that we did not observe a significant difference in in synaptic density and excitatory pre-synaptic puncta between the tri- and co-cultures may be due to the fact that the microglia and neurons are in close approximation in the tri-culture leading to a balance of increased synaptogenesis due to IL-10 secretion by microglia and synaptic pruning due to the direct contact of microglia and synapses. Furthermore, astrocytes have also been shown to play a significant role in synaptic plasticity [62], and since both the tri- and co-cultures contain astrocytes, the influence of the microglia may be less apparent.

We also investigated the E/I ratio as a potential cause of the increased spike frequency and found that we recorded from a significantly higher percentage of broad-spiking excitatory neurons in the tri-culture (**Figure 4b**). Additionally, the E/I ratio of the tri-culture was much closer to the E/I ratio found *in vivo* [50]. However, we cannot be sure if this apparent increase in excitatory neurons in the tri-culture is due to an actual increase in the number of excitatory neurons in the tri-culture or due to a relative increase in the number of excitatory neurons incorporated into the neural network. It has been shown that GABAergic neurons may be less healthy than glutamatergic neurons in culture [63], and it may be possible that the microglia in the tri-culture are inducing apoptosis in the weakened GABAergic neurons. Alternatively, it has also been shown that glutamatergic neurons are much more dependent on glial support to establish functional glutamatergic synapses [63], and therefore the presence of microglia in the tri-culture may further enhance the incorporation of excitatory neurons into the neural network. Additionally, microglia are known to regulate perineuronal nets (PNNs), which stabilize synapses and have been shown to appear in cortical neuron cultures [64]. These PNNs primarily associate with narrow-spiking inhibitory neurons, leading to their over incorporation into neural networks when microglia are not present [43]. Since microglia are present in the tri-culture, they may be able to regulate the PNNs, thereby allowing increased incorporation of excitatory neurons into the neural network and increasing the apparent E/I ratio to values closer to what is observed *in vivo*.

Finally, we investigated the tri-culture’s ability to detect changes in neural activity in response to neuroinflammatory stimuli. We therefore challenged the tri- and co-culture with 5 μg/mL LPS and monitored the change in neural activity over 72 hours. LPS is a well characterized activator of neurotoxic neuroinflammation that acts through the TLR4, which is found on microglia, but not neurons and astrocytes [65,66]. Activation of microglia by LPS leads to the secretion of proinflammatory cytokines [32,67], neuronal apoptosis [68,69], and the induction of a neurotoxic “A1” astrocyte phenotype [70]. Additionally, we have demonstrated a significant increase in apoptosis and cell death in tri-cultures treated with 5 μg/mL LPS at 48 h [32,33]. Consequently, we observed a sharp decrease in both the number of active channels and spike frequency in the LPS treated tri-cultures that became significant by 12 h post exposure (**Figure 5**). Additionally, we observed that the spike frequency decreases at a more rapid rate than the number of active channels, suggesting that changes in neural activity can be detected prior to changes in neuron health. This observation can be paralleled to the findings from the recent study that showed a decrease in electrophysiological activity with increasing microglia density in a dose-response manner [54], where higher microglia numbers in the culture lead to increased microglia reactivity, partially mimicking increased reactivity due to the LPS treatment here. This demonstrates that monitoring extracellular recordings from MEAs is a powerful method to non-invasively study the effect of neuroinflammation *in vitro*, especially when combined with our previously described tri-culture of neurons, astrocytes, and microglia that more faithfully represents the *in vivo* neuroinflammatory response.

## Conclusion

In this study, we characterized the neural activity and network maturation in a tri-culture of neurons, astrocytes, and microglia. We showed that the presence of microglia in the tri-culture had minimal impact on the formation and stability of neural networks, with the exception of an increase in spike frequency in the tri-culture. Additionally, characterization of the average spike waveforms revealed that the tri-culture had an E/I ratio much closer to that found in the rat cortex. Finally, we demonstrated that the more biologically-relevant neuroinflammatory response of the tri-culture can be captured via extracellular recordings as indicated by a significant decrease in both active channels and spike frequency following exposure to LPS. We expect that the electrophysiological read-out from the tri-culture will be useful for continuous and non-invasive studies in the context of neuroinflammation, neurodevelopment, and neurodegeneration, where the presence of microglia imparts the ability to capture both neurotoxic and neuroprotective phenomena observed *in vivo*.

## Abbreviations

MEA: Microelectrode array
E/I ratio: excitatory to inhibitory neuron ratio
LPS: Lipopolysaccharide
CNS: Central nervous system
TLR: Toll-like receptor
DIV: Days in vitro
PDMS: Polydimethylsiloxane
PECVD: Plasma enhanced chemical vapor deposition
IL: Interleukin
TGF-β: Transforming growth factor beta
DPBS+: Dulbecco’s phosphate buffered saline solution with calcium and magnesium
PFA: Paraformaldehyde
PSD-95: Postsynaptic density protein 95
VGlut1: Vesicular glutamate transporter 1
MAP2: microtubule associated protein 2
GFAP: Glial fibrillary acidic protein
Iba-1: Ionized calcium binding adaptor molecule 1
DAPI: 4’,6-diamidino-2-phenylindole
SEM: Standard error of the mean
ANOVA: Analysis of variance
TLR4: Toll-like receptor 4
PNN: Perineuronal nets

## Ethics Approval and Consent to Participate

All procedures involving animals were conducted in accordance with the National Institutes of Health Guide for the Care and Use of Laboratory Animals following protocols approved by the University of California, Davis Institutional Animal Care and Use Committee.

## Consent for Publication

Not applicable.

## Availability of Data and Materials

he datasets during and/or analyzed during the current study are available from the corresponding author on reasonable request.

## Author Contributions

NG and ES designed the experiments. NG performed device fabrication, cell culture, electrophysiological recordings, imaging, and data analysis and wrote the main manuscript text. GG contributed to the development and fabrication of microfluidic devices and microelectrode arrays, and data analysis. HK conducted confocal microscopy. AG contributed to electrophysiological data analysis. ES contributed to the interpretation of experimental results and edited the manuscript. All authors read and approved the final manuscript.

## Acknowledgements

This project benefited from the resources of the MIND Institute IDDRC Core services (NICHD P50-HD103526) and University of California, Davis - Center for Nano/Micro-Manufacturing facility.

