## Supplemental Information for "Electrophysiological Activity of Primary Cortical Neuron-Glia Mixed Cultures"

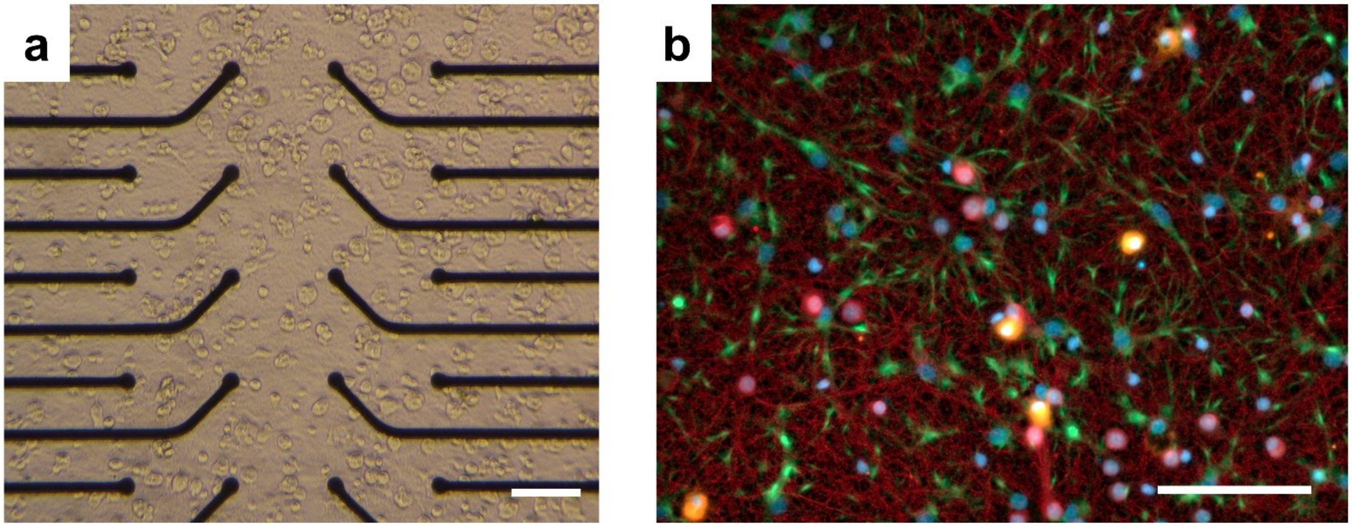


**Figure S1:** (a) Bright-field and (b) epifluorescence images of the tri-culture at DIV 21 on the well MEA. The cultures were immunostained for the three cell types of interest: neurons—anti-βIII-tubulin (red), astrocytes—anti-GFAP (green), microglia—anti-Iba1 (orange), and the general nuclear stain DAPI (blue). Scale bar = 100 μM.

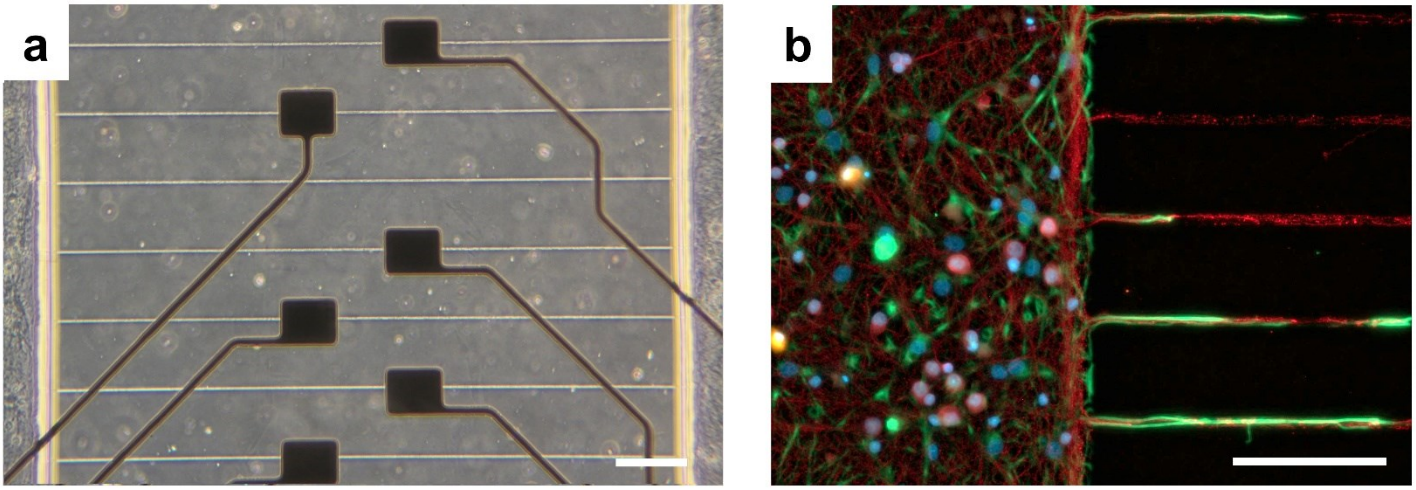


**Figure S2:** (a) Bright-field and (b) fluorescence images of the tri-culture at DIV 21 in the platform MEA with microfluidic encapsulation. The cultures were immunostained for the three cell types of interest: neurons—anti-βIII-tubulin (red), astrocytes—anti-GFAP (green), microglia—anti-Iba1 (orange), and the general nuclear stain DAPI (blue). Scale bar = 100 μM.

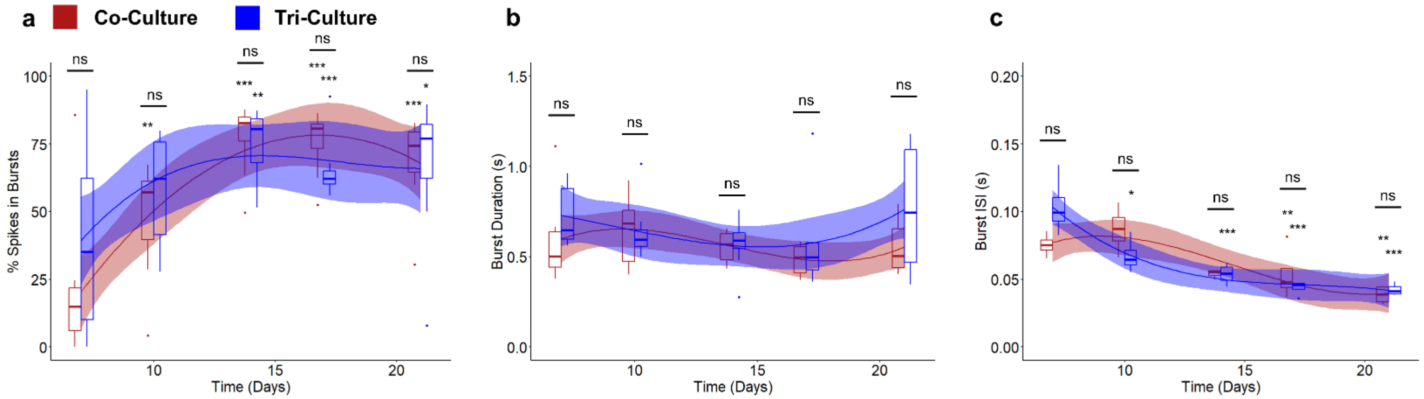


**Figure S3:** Comparisons of the (a) percent spikes in bursts, (d) burst duration, and (e) ISI within bursts between co-cultures (red) and tri-cultures (blue). The solid lines show the fitted linear mixed effects model (treating individual cultures as a random effect) with a b-spline basis. The shaded regions are the 95% confidence interval. An asterisk above an individual box indicates a significant difference of the estimated marginal means of the fitted curves between that timepoint and DIV 7 of the same culture type, while the bars indicate the significance between the co- and tri-culture at that timepoint (n = 8, from three independent dissections). *$p < 0.05$, **$p < 0.01$, ***$p < 0.001$, ns indicates no significant difference.

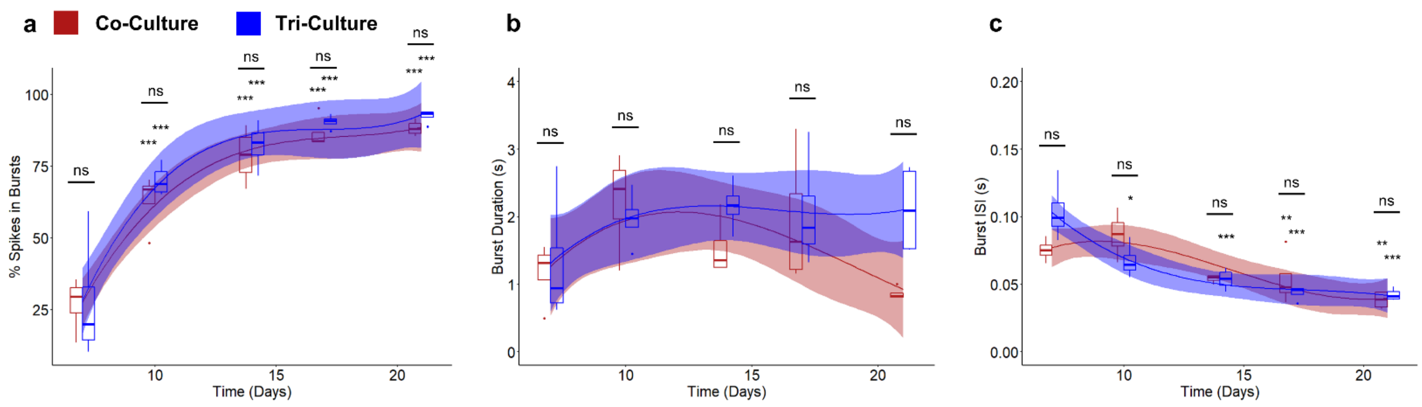


**Figure S4:** Comparisons of the (a) percent spikes in bursts, (d) burst duration, and (e) ISI within bursts between co-cultures (red) and tri-cultures (blue) cultured in a two chambered microfluidic device. The solid lines show the fitted linear mixed effects model (treating individual cultures as a random effect) with a b-spline basis. The shaded regions are the 95% confidence interval. An asterisk above an individual box indicates a significant difference of the estimated marginal means of the fitted curves between that timepoint and DIV 7 of the same culture type, while the bars indicate the significance between the co- and tri-culture at that timepoint (n = 5, from two independent dissections). *$p < 0.05$, **$p < 0.01$, ***$p < 0.001$, ns indicates no significant difference.

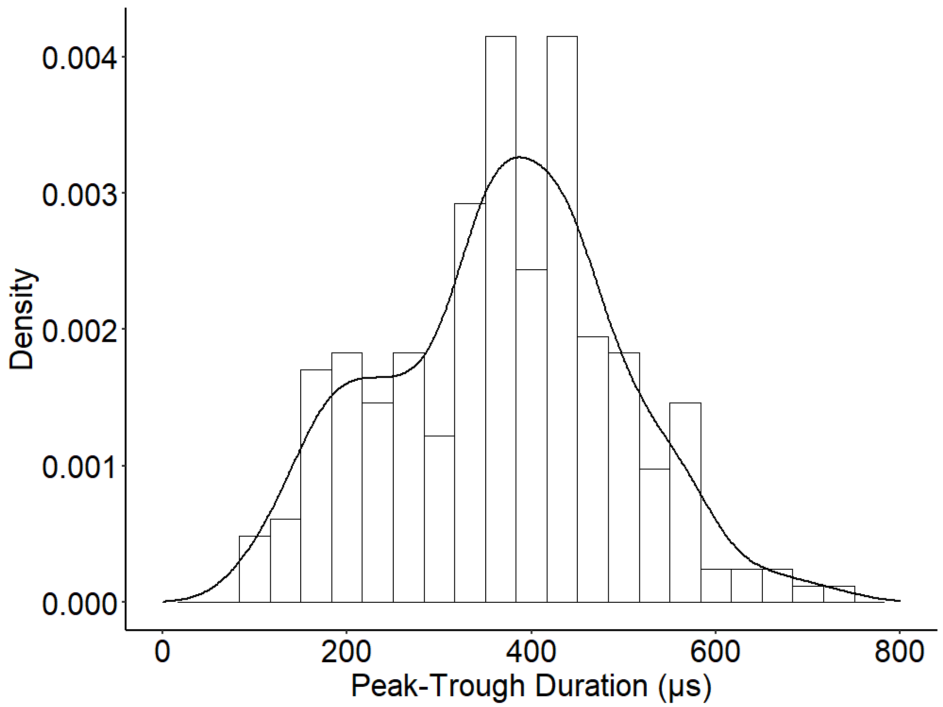


**Figure S5:** Density histogram of the peak-trough durations of the averaged spike waveforms from 249 units recorded from both co- and tri-cultures. We observe one peak at ~220 µs and the second at ~380 µs.
